# Quantifying the risk of hemiplasy in phylogenetic inference

**DOI:** 10.1101/391391

**Authors:** Rafael F. Guerrero, Matthew W. Hahn

## Abstract

Convergent evolution is often inferred when a trait is incongruent with the species tree. However, trait incongruence can also arise from changes that occur on discordant gene trees, a process referred to as hemiplasy. Hemiplasy is rarely taken into account in studies of convergent evolution, despite the fact that phylogenomic studies have revealed rampant discordance. Here, we study the relative probabilities of homoplasy (including convergence and reversal) and hemiplasy for an incongruent trait. We derive expressions for the probabilities of the two events, showing that they depend on many of the same parameters. We find that hemiplasy is as likely— or more likely—than homoplasy for a wide range of conditions, even when levels of discordance are low. We also present a new method to calculate the ratio of these two probabilities (the “hemiplasy risk factor”) along the branches of a phylogeny of arbitrary length. Such calculations can be applied to any tree in order to identify when and where incongruent traits may be more likely to be due to hemiplasy than homoplasy.

## Introduction

Convergent traits found in distantly related organisms are prime examples of the role of natural selection in evolution. They are often used as evidence for the importance of adaptation in shaping organismal form and function, though they may also reflect underlying constraints on developmental pathways (1). Understanding how often convergent evolution occurs, and the conditions under which it occurs, will allow us to better understand its causes. The identification of clear cases of convergence—or more broadly, homoplasy, which includes reversals to ancestral states (2)—also enables us to determine how often convergent phenotypes are underlain by convergent molecular changes (3, 4).

Homoplasy, whether due to convergence or reversal, is inferred when multiple evolutionary changes are required to explain the character states observed among sampled lineages.

Therefore, in order to make strong inferences about homoplasy we require both a phylogenetic tree describing the relationships among taxa and a model of trait evolution. If either the tree or the model is incorrect, this can lead to errors in inferences about the number of transitions that have occurred (5–9). In order to reduce errors in species trees, genome-scale datasets have been used to generate topologies that have strong statistical support at almost all nodes (e.g. (10, 11)). Though concerns remain about appropriate models of trait evolution in some cases (12), the “resolution” of species trees with phylogenomic datasets would appear to have removed the main source of error in inferring character-state transitions accurately.

However, genome-wide data have also highlighted the ubiquity of gene tree discordance, even when statistical support for the species tree is high (e.g. (13–16)). Discordance can be due to many factors, both technical and biological. The major biological causes of gene tree discordance are incomplete lineage sorting (ILS) and introgression (17). In the presence of either of these processes, individual loci can have different histories from the species tree; discordance due to either introgression or ILS does not go away over time, so studies of both ancient and recent divergences can be affected.

Discordance presents a problem for inferences of convergent evolution because it can produce patterns of homoplasy even when none has occurred (18). In a phenomenon dubbed “hemiplasy” (19), transitions that occur on branches of discordant trees that do not exist in the species tree will generate incongruent trait patterns (Figure 1A). Incongruent trait patterns— those that cannot be explained by a single trait transition—are the basis for claims of convergent evolution and homoplasy. As all discordant trees have discordant branches (20, 21), hemiplasy is expected to explain a substantial number of observed incongruent trait patterns in all cases where biological discordance exists (22–25).

**Figure 1.**
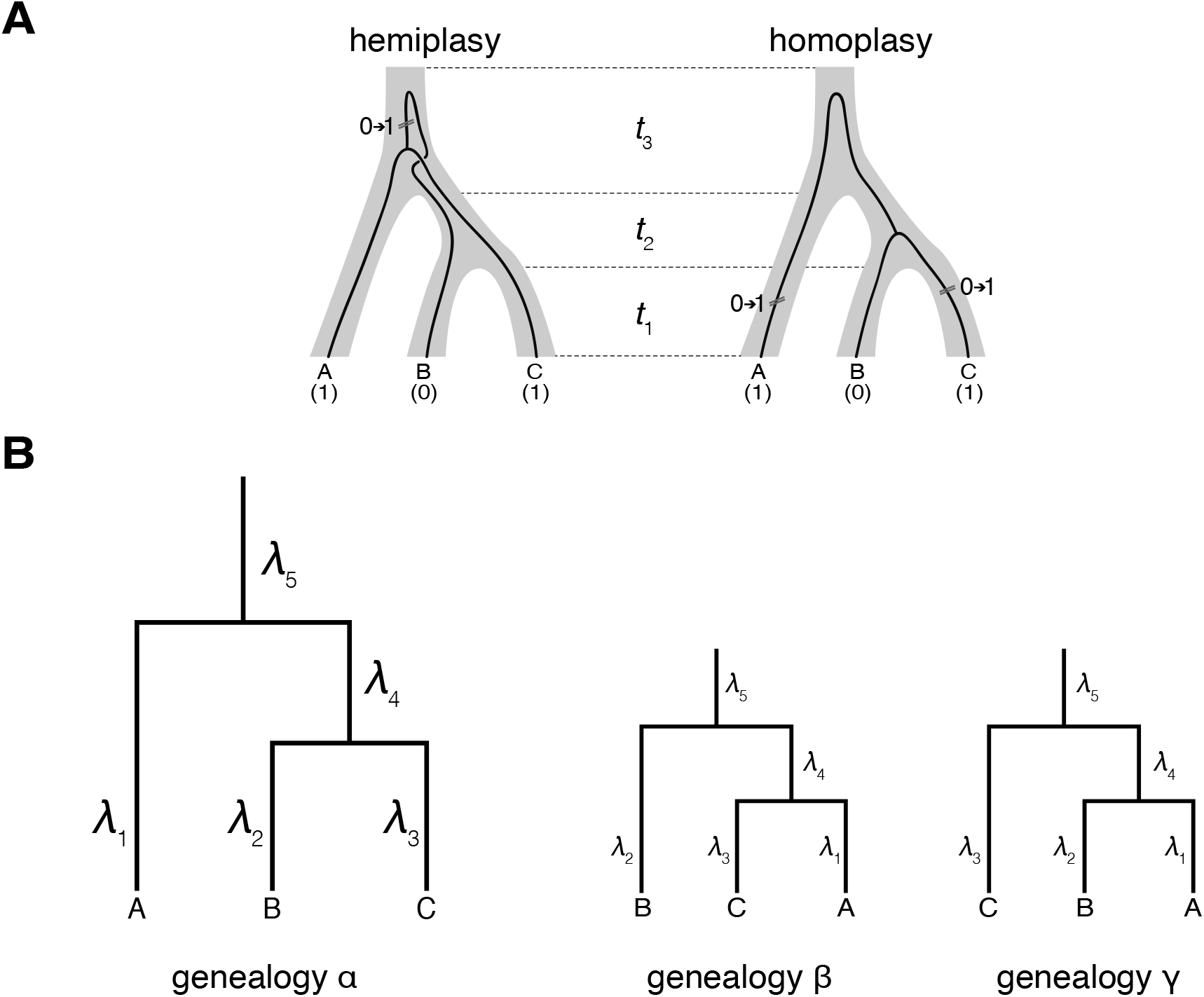
A) The same species tree is shown for both hemiplasy and homoplasy, with times in this tree denoted with *t_i_* labels. Left: an example of how hemiplasy can generate an incongruent trait pattern among species A, B, and C. A single mutation on the internal branch of a discordant gene tree (denoted as 0→1) produces this pattern. Right: an example of how homoplasy (via convergence) can generate the same incongruent trait pattern via two mutations from 0→1. B) Branch labeling for the genealogies α, β, and γ.

While complications due to hemiplasy have begun to be appreciated, the relative importance of hemiplasy and homoplasy in any particular set of relationships has yet to be quantified. Here, we present a model that describes the relative probabilities of hemiplasy and homoplasy for an incongruent trait. Given even minimal amounts of gene tree discordance, we find that hemiplasy is more likely, or at least as likely, as homoplasy for a wide range of conditions. Based on this model, we present a method for calculating the ratio of hemiplasy to homoplasy (the “hemiplasy risk factor”) along a phylogeny. This method provides a general tool for understanding the risk of incorrect inferences of convergence, for use with phylogenies of any size.

## Results

### The model

Consider the evolution of a trait among three species with the phylogeny (A,(B,C)). We assume that this trait, which can be either molecular or phenotypic, is binary with 0 representing the ancestral state and 1 representing the derived state. On this phylogeny we observe an incongruent trait pattern (e.g., allelic states of A=1, B=0, and C=1; as in Figure 1A) that cannot be explained by a single transition along the species tree. There are two possible explanations for this incongruence (Figure 1A). One possibility is that the trait is hemiplastic: a single transition has occurred along the internal branch of a discordant gene tree in which B is sister to a clade consisting of A and C. Alternatively, the trait is homoplastic: two trait transitions have occurred, due to either convergence (two changes from 0→1) or reversal (one change from 0→1 and one change from 1→0; Supplementary Figure 1). In both homoplasy scenarios, some of the observed allelic states are not identical by descent, though they differ as to whether these are the derived states (convergence) or the ancestral states (reversal).

The goal of our model is to understand the biological parameters that affect the relative frequency of these two mechanisms for trait incongruence. To address this goal, we derive expressions for the probabilities of hemiplasy (*P_e_*) and homoplasy (*P_o_*), which both broadly depend on population size (*N* diploid individuals, assumed constant), the rate of mutation (μ, per 2*N* generations), and the timing of speciation events (in units of 2*N* generations). Assuming that all discordance is due to incomplete lineage sorting, the population size and timing of speciation events determine the genealogy τ, with possible topologies α=(A,(B,C)), β=(B,(A,C)), and γ=(C,(A,B)). Topology α is concordant with the species tree, while both β and γ are discordant. Classic results from coalescent theory (26) provide the probability of each topology, along with the lengths of each branch in each tree. The topology of tree τ determines the probability of mutation, ν(λ_*i*_,τ), along every branch *i* (labeled as in Figure 1B) with length λ_*i*_. Because of its dependence on gene tree topology, the function ν(λ_*i*_,τ) is different for each branch. For instance, the probability of mutation along the internal branch in discordant tree β is uniquely constrained by *t*_3_ and the time to coalescence of A and C. The expressions for each ν(λ_*i*_,τ) are described in the Methods.

In the case of hemiplasy, two events are necessary to explain the allelic states specified above: the occurrence of discordant genealogy β and a mutation occurring on the internal branch of this gene tree (but not on any other branch). This probability is given by:

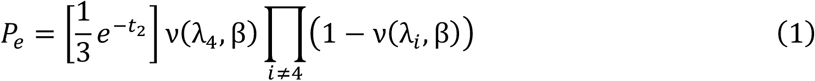

where *t_2_* is the length of the single internal branch on the species tree (Figure 1A), and λ_4_ is the length of the internal branch on genealogy β (Figure 1B). The three terms here correspond to (from left to right): the probability of observing genealogy β, the probability of a mutation occurring on the internal branch of this genealogy, and the probability of no mutations occurring anywhere else.

In the case of homoplasy, two alternative sets of mutational events may have occurred: either two independent origins of the derived state (on the terminal branches leading to A and C), or a single mutation along the branch leading to the ancestor of all three species, coupled with a back-mutation on the branch leading to B. Because these events can occur on any one of the three possible topologies, the probability of homoplasy is:

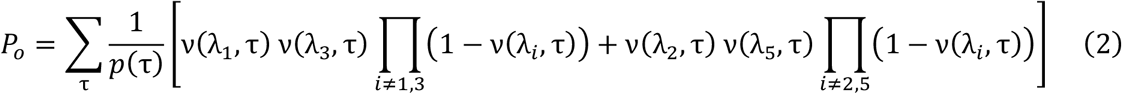

where ***p***(***τ***) is the probability of genealogy τ. The probability of homoplasy therefore corresponds to the sum of the probability of convergence averaged across all topologies and the probability of reversal averaged across all topologies (the first and second terms in the summation, respectively).

### The relative probability of hemiplasy and homoplasy

Our model demonstrates that the probabilities of both hemiplasy and homoplasy are determined by many of the same factors. The frequencies of individual gene trees, the branch lengths in these trees, and the rate of mutation to the character of interest all interact to determine these probabilities. However, because the exact values for these parameters may rarely be known, the precise probabilities of each of these outcomes on their own may not be easily obtainable. Instead, here we present the ratio of probabilities of hemiplasy and homoplasy (*P_e_/P_o_*) for a range of parameter values in order to provide a quantitative sense for when each will be relatively more important.

The amount of discordance (which decreases with increasing *t_2_*) and the mutation rate tend to have opposite effects on the relative probabilities of hemiplasy and homoplasy (Figure 2A).

**Figure 2.**
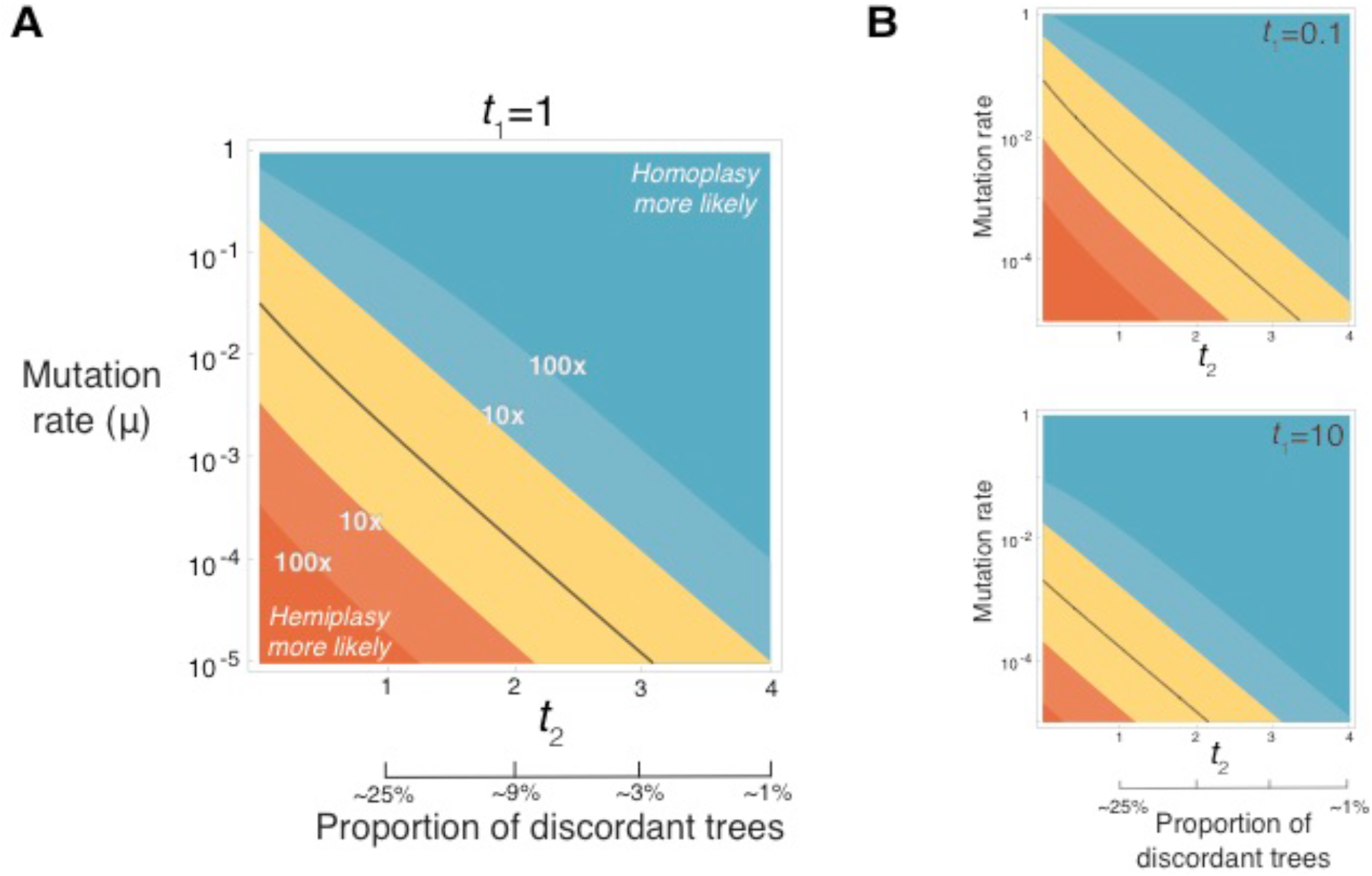
A) The relative probabilities of hemiplasy and homoplasy (*P_e_/P_o_*) for a range of values of *t*_2_ (in units of 2*N* generations) and the rate of mutation (μ, per 2*N* generations). The black solid line represents parameter values for which hemiplasy and homoplasy are exactly equal (*P_e_/P_o_* =1). We also show the expected proportion of discordant trees for various values of *t*_2_ on the x-axis. The values of *t*_1_=1 and *t*_3_=4 are fixed. B) The effect of changing *t*_1_ on *P_e_/P_o_*. The value of *t*_3_ remains the same as in panel A.

As expected, increasing levels of discordance lead to increasing relative probabilities of hemiplasy. Although homoplasy can occur on discordant trees, this outcome still requires two mutations; therefore, for a given mutation rate, more discordance will always lead to relatively more hemiplasy than homoplasy. On the other hand, when there is little opportunity for discordance, there is also a greatly reduced probability for hemiplasy. By the time an internal branch of a species tree is 6*N* generations long, approximately 99.9% of all gene tree topologies will share this branch—that is, they will be concordant. In this area of parameter space homoplasy becomes vastly more likely than hemiplasy as an explanation for incongruent trait patterns.

The mutation rate also has a strong effect on the relative probabilities of hemiplasy and homoplasy. As should be expected, when the rate of mutation to the character of interest is high, homoplasy becomes relatively more likely. Reducing mutation rates results in a steeper decrease in homoplasy relative to hemiplasy (because of the two mutations that are required), leading to a higher ratio of *P_e_/P_o_*. For low enough mutation rates, very few trees must be discordant in order for the probabilities of hemiplasy and homoplasy to be equal, so that *P_e_/P_o_*=1. In this area of parameter space there is an equal probability for any particular incongruent trait pattern to be caused by either hemiplasy or homoplasy. We can also think of this ratio as indicating that, given a collection of incongruent trait patterns of this type, we expect approximately half will be due to hemiplasy and half to homoplasy. These results, shown for one particular pattern of incongruence (A=1, B=0, C=1), are expected to be the same for the two possible incongruence patterns on the species tree in Figure 1A (see the next section for cases in which these can be different).

While the amount of discordance and rate of mutation both have strong effects on *P_e_/P_o_*, the length of the branch subtending the clade of interest (*t_3_*; Figure 1A) and the lengths of the tip branches (*t_1_*) have a much smaller effect. The length of *t_3_* is only relevant to cases of homoplasy caused by reversal and has no effect on either the probability of hemiplasy or the probability of homoplasy caused by convergence. As such, it has very little effect on the relative probabilities of these two outcomes (Supplementary Figure 2). In contrast, *t_1_* affects the probability of homoplasy by both reversal and convergence while having no impact on the probability of hemiplasy, and consequently has a larger influence on *P_e_/P_o_* (Figure 2B). All convergent events on the species tree considered here involve changes on one of the paired lineages (in this case species C) and the unpaired lineage (species A), so *t_1_* determines how much time there is for a change to occur on these lineages (together with *t_2_*, which contributes to the probability of a mutation along the branch leading to species A). Therefore, larger or smaller values of *t_1_* allow for more or less time in which homoplasy can occur, respectively, shifting the balance of *P_e_/P_o_* slightly (compare Figure 2A to 2B).

A numerical example may help to highlight the utility of these calculations. One of the most well-studied cases of discordance caused by incomplete lineage sorting occurs among human (H), chimpanzee (C), and gorilla (G; briefly reviewed in (27)). The genome sequences of these three species—plus an outgroup (orangutan)—show that 70% of nucleotides in protein-coding regions are congruent with the presumed species tree (i.e., (G,(H,C))), with 15% congruent with each of the (C,(H,G)) and (H,(C,G)) topologies (28). Multiple studies have also calculated the divergence times and effective population sizes of these species and their ancestral lineages (e.g. (29, 30)), as well as per-generation nucleotide mutation rates (e.g. (31)), offering us the opportunity to estimate *P_e_/P_o_* for this clade. For incongruent nucleotides with states H=1, C=0, and G=1, the expected *P_e_/P_o_* = 5.4 (see Methods). This estimate means that hemiplasy is roughly five times more likely than homoplasy to explain cases where human and gorilla share a derived allele to the exclusion of chimpanzee. Alternatively, we can view the value as indicating that hemiplasy is the cause of approximately 84% of all sites with this incongruent site pattern. This value of *P_e_/P_o_* uses the average mutation rate per site in the genome (1.20 x 10^−8^ per generation), even though the rate can vary by orders of magnitude across sites. At hypermutable CpGs (1.22 x 10^−7^; (31)), for instance, only one-third of observed incongruence is expected to be due to hemiplasy (i.e., *P_e_/P_o_* = 0.52). In contrast, at non-CpG sites (9.94 x 10^−9^; (31)) approximately 87% of all incongruence will be due to hemiplasy (i.e., *P_e_/P_o_* = 6.5)

### Calculating the hemiplasy risk factor along a phylogeny

The model and results presented thus far have focused on a rooted three-taxon tree, with speciation events that have occurred in the recent past. Importantly, however, the relevant calculations for the probabilities of hemiplasy and homoplasy are not restricted to such trees.

They can be applied to trees of any size, of any height, and with non-ultrametric branch lengths.

These probabilities can therefore be used with larger phylogenetic trees to highlight individual branches along which hemiplasy is likely to be responsible for observed incongruence.

In order to usefully summarize the relative probabilities of hemiplasy and homoplasy, we define the “hemiplasy risk factor” (HRF) as the ratio of *P_e_* to *P_o_* averaged across incongruent trait patterns:

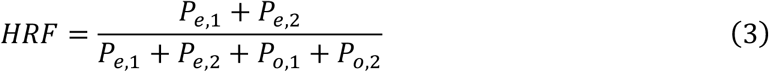

where the subscripts “1” and “2” represent the two possible incongruent trait patterns among three species. Although the probabilities of hemiplasy and homoplasy can be found for phylogenetic “knots” of more than three lineages (cf. (20, 32)), we confine HRFs to the three-taxon case so that we can efficiently calculate them across larger species trees (see below). We calculate the values of *P_e_* and *P_o_* separately for each incongruent pattern because in non-ultrametric trees these may be quite different from one another (e.g. when the branches leading to species “B” and “C” are not the same length; Figure 1A). While this means that the HRF no longer represents the probability of hemiplasy for a specific incongruent pattern, it does helpfully summarize the overall risk of hemiplasy relative to homoplasy.

The HRF is intended to highlight individual branches of phylogenetic trees along which hemiplasy may be responsible for observed incongruence, in opposition to the assumption that all incongruence is due to homoplasy. An HRF can be calculated for each internal branch of a rooted species tree, but not for tip branches. Note that given the definitions of hemiplasy and homoplasy, the HRF associated with a branch does not indicate that a character-state transition has occurred along this branch: for instance, under hemiplasy due to ILS the relevant mutation has occurred on an earlier lineage (e.g. the one with length *t_3_* in Figure 1A) but has remained polymorphic through the relevant branch (e.g. the one with length *t_2_* in Figure 1A). Instead, HRFs identify branches of a tree where the processes leading to hemiplasy may be occurring, and around which homoplastic transitions may be incorrectly inferred. In such cases standard ancestral state reconstruction methods will infer homoplastic substitutions on the branches neighboring the one with the high HRF—either the branches directly “above” and “below” it in the case of reversals, or one branch below and one branch sister to it in the case of convergence (Supplementary Figure 1).

We have implemented a package written in R (www.r-project.org) for calculating and visualizing HRFs on larger phylogenetic trees (github.com/guerreror/pepo). The software walks through a phylogenetic tree starting from the tips, calculating HRFs on every trio of lineages (Methods). The input species tree must have branch lengths given in units of 2*N* generations (sometimes referred to as “coalescent units”), and a mutation rate must be specified. Multiple methods can output species trees in coalescent units (e.g. MP-EST; (33)), or—under the assumption that all gene tree discordance is due to ILS—these lengths can be estimated for internal branches of a tree by taking the proportion of discordant trees (i.e. 1 minus the concordance factor; CF) and solving for *t_2_* = 3/2 ln(1 − *CF*). Because HRFs are intended to aid researchers during exploratory studies of the evolution of many different types of traits, the exact value of the mutation rate used should not be a key concern. For a reasonable estimate of the mutation rate the HRFs calculated will instead represent the relative risk among branches of a larger tree along which hemiplasy may be occurring. The exact value of *P_e_/P_o_* can still be calculated for a specific trait of interest on a smaller portion of the tree using equations 1 and 2.

To provide an example of HRFs calculated on a larger tree, we use the phylogeny of wild tomatoes presented in Pease et al. (15). This dataset represents a rather extreme example of a recent rapid radiation, with none of the 2745 gene trees (actually inferred from 100-kb windows) matching the exact topology of the species tree. After generating the necessary input species tree from a collection of gene trees (Methods), we calculated HRFs for all internal branches (Figure 3). As can be seen, while there are multiple longer branches along which homoplasy should be a better explanation for incongruent patterns than hemiplasy, there are also many branches with equal or higher relative probabilities of hemiplasy. The branches with high HRFs are often the shortest branches, where the most discordance is to be expected. But note that two branches with equal discordance (i.e. equal concordance factors; (32)) can have very different HRFs. The calculation of HRFs depends not only on the length of the target branch in coalescent units— which solely determines the expected degree of discordance, and to a large extent the probability of hemiplasy—but also on the length of the surrounding branches, which help to determine the probability of homoplasy. HRFs therefore represent a unique and complementary tool for understanding trait evolution on trees.

**Figure 3.**
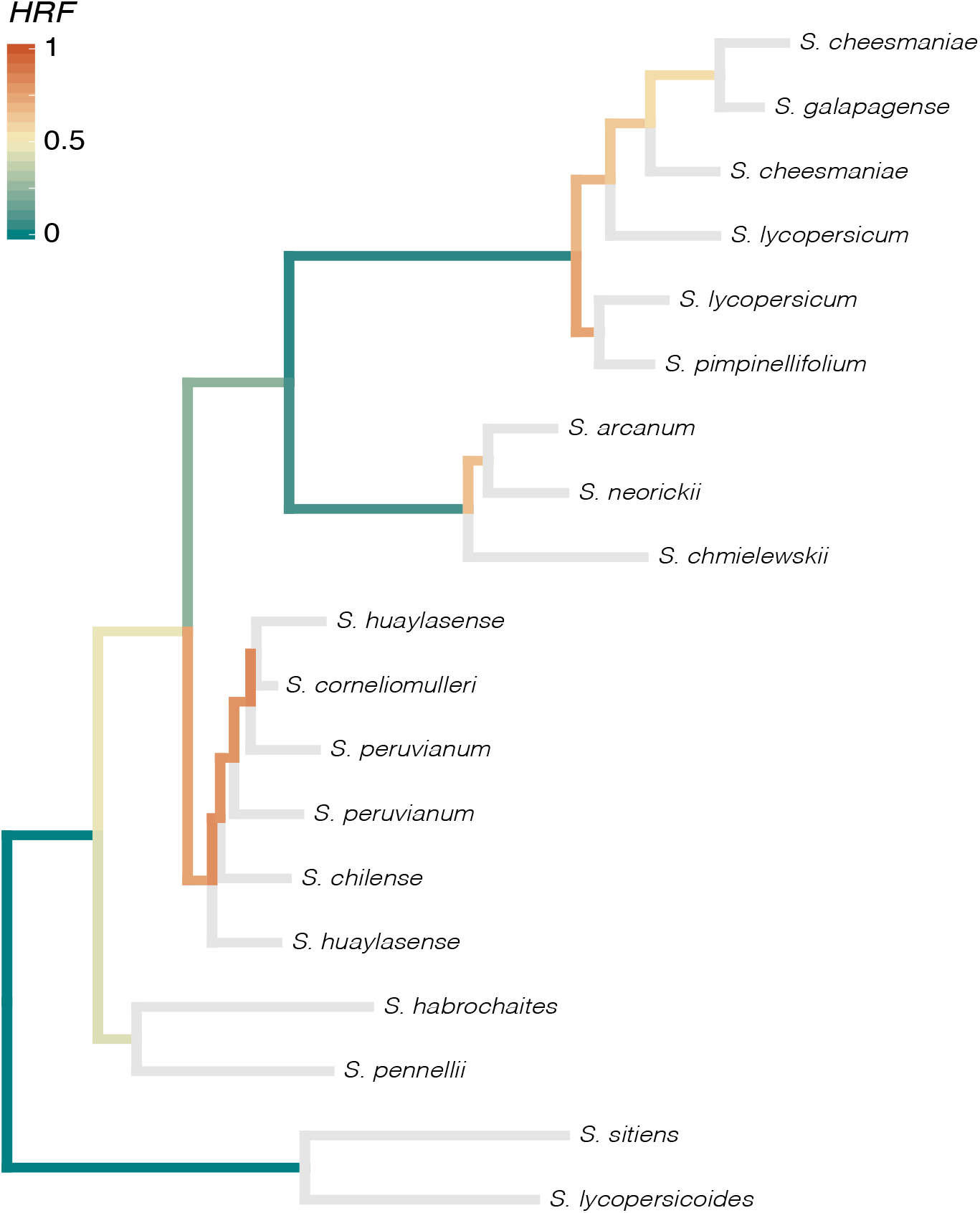
Inferred tree for species in the genus *Solanum* labeled with hemiplasy risk factors (HRFs). The species tree (with branch lengths in units of 2*N* generations) was inferred using gene trees from (15); see Methods for details. HRFs were calculated for all internal branches assuming μ=0.01 per 2*N* generations. Values represent the proportion of incongruent traits associated with branch that are due to hemiplasy, so that HRF=0 means no hemiplasy and HRF=1 means all patterns due to hemiplasy.

## Discussion

Phylogenomic studies have revealed high levels of gene tree discordance in many species trees (13–16). Such discordance can be concealed by statistical measures of support for the species tree, such as bootstrap support or posterior probabilities, which can be high even when discordance is rampant. Regardless of confidence in the species tree topology, underlying gene tree discordance means that observed patterns of trait incongruence can be due to single transitions. This phenomenon of hemiplasy likely explains the distribution of multiple ecologically important traits (e.g. (34, 35)), but can mislead standard methods for inferring the number of times a trait has involved and the timing of these transitions (23–25). The goal of the work presented here is to quantify the risk of incorrectly inferring homoplasy when hemiplasy is occurring. We do this by finding explicit expressions for the probabilities of hemiplasy and homoplasy, and use these expressions to develop a new measure (the hemiplasy risk factor) that highlights branches of a phylogeny along which hemiplasy may be occurring.

Our model is currently applicable to any type of binary trait, as these are the traits that are most often the focus of studies of convergence. While it should be straightforward to extend the model to any sort of discrete trait (e.g. gene family size (36)), even when limited to binary traits a more important question may be what value of the mutation rate to use. We have defined the mutation rate in our model as the rate at which one state can change into the other. Although we have discussed a reversible process here (i.e. mutations from 0→1 and 1→0 are allowed), this is not a requirement of our model—it can accommodate other types of mutational processes (e.g. (37)). We do not need to specify whether the trait is molecular or phenotypic, nor any other details about the process. For studies of molecular traits—such as nucleotide substitutions—the appropriate mutation rate to use should be clear. For phenotypic traits the mutation rate may be much higher or much lower than this per-nucleotide rate. For example, the loss of some traits may be possible by the inactivation of multiple genes (e.g. (38)); in such cases the appropriate mutation rate should include the sum of nucleotide mutation probabilities at all the sites in the genome at which inactivation can be accomplished, making it much higher than the single-site rate. In contrast, trait transitions requiring multiple nucleotide changes at a limited number of sites in the genome may have a total mutation rate that is equal to the square or cube of the per-nucleotide rate, making it even lower (and consequently making homoplasy even less likely). Although the hemiplasy risk factor still requires a mutation rate be specified in order to be calculated, our hope is that it will highlight the relative risk among different branches of a larger tree regardless of the specific value used.

Our model makes two important assumptions. First, we have based all of our calculations on a coalescent model in which ILS is the only process that generates gene tree discordance. However, at least among eukaryotic organisms, introgression appears to be a cause of discordance in a wide variety of systems (39). To some extent it should not matter what process causes discordance, as both ILS and introgression will be more likely to lead to discordant topologies when there are short internode branches (because introgression between sister lineages does not result in discordance). The hemiplasy risk factor will therefore still point to the same lineages along which hemiplasy is high or low relative to homoplasy, the latter of which is not affected by introgression. In addition, our calculations assuming only ILS are likely to be underestimates of the probability of hemiplasy: if the cause of discordance is introgression, the internal branches of discordant gene trees do not have to be as short, making the probability of mutation along them higher.

Our second important assumption is that selection is not acting on the traits of interest. Directional selection on loci underlying a trait will cause them to be more concordant relative to the neutral expectations for the case in which ILS is the sole cause of discordance. Note, however, that the magnitude of this effect may be small, especially for traits controlled by multiple loci. Even when selection on individual loci is strong, it is only selection on the internal branch of the species tree that matters—lineage-specific selection cannot affect discordance. Moreover, if introgression is the cause of hemiplasy, then selection is irrelevant to the probability of discordance. While directional selection decreases discordance, balancing selection can increase the probability of discordance. Multiple examples of hemiplasy acting on balanced polymorphisms have already been identified (e.g. (13, 34, 40)). As these examples are also associated with shorter internal branches of the respective species trees, the hemiplasy risk factor will likely also have highlighted the pertinent branches, regardless of whether the assumptions of our model have been violated.

Recent work on the genetic basis of convergent traits has revealed that such traits are sometimes determined by convergent molecular changes (e.g. (41–43)). The proportion of cases in which convergent phenotypes are underlain by convergent genotypes remains an open question (3, 4), and both additional phylogenetic and functional work will be needed to accurately estimate this proportion. The work here aims to aid phylogenetic studies of convergent evolution by quantifying the proportion of time the apparent convergence in traits (which would also appear to be molecular convergence) may instead be caused by underlying gene tree discordance. Although the ecological conditions under which hemiplasy occurs may still be informative about the processes driving similarity in traits (e.g. (44)), properly distinguishing between hemiplasy and homoplasy is necessary for understanding the molecular basis for convergent evolution.

## Methods

### Probabilities of mutation along genealogies

To clarify the calculations of *P_e_* and *P_o_* in equations 1 and 2, we must specify how the probability of mutation is calculated for each branch. On random branch lengths (as in the case of coalescent times), the probability of mutation is 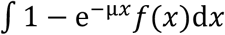, where *f*(*x*) is the probability density function of *x* (the random variable for branch length). All mutation probabilities ν(λ_*i*_,τ) have some version of that general form, varying the value of *x* and *f*(*x*). Genealogies β and γ can only happen in the absence of coalescence between lineages B and C in the BC ancestor. In contrast, genealogy α can happen with or without coalescence in BC, and it is helpful to consider the two alternatives separately: α_0_ denotes the subset of α trees that coalesce in the ABC ancestor (i.e., without coalescence in BC), whereas α_+_ are the trees that have the α topology and coalesced in BC.

The genealogies α_0_, β, and γ are identical in length (although they have different tip identities), and their mutation probabilities can be described by the equations:

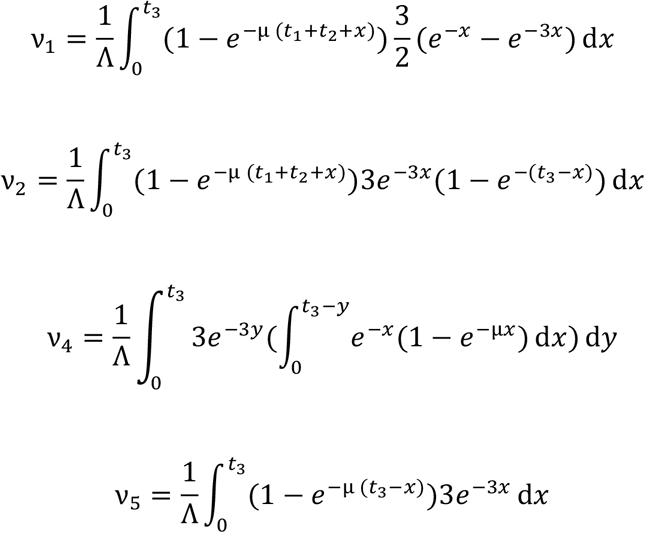

In the above, 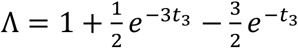 and represents the probability of coalescence of A, B, and C in the ABC ancestor (i.e., the cumulative distribution function of the coalescent for a sample of size 3). Each of these four probabilities correspond to multiple branches in Figure 1B. Specifically, ν_1_ = ν(λ_1_, α_0_) = ν(λ_2_, β) = ν(λ_3_,γ), ν_2_ = ν(λ_2_, α_0_) = ν(λ_3_, α_0_) = ν(λ_1_, β) = ν(λ_3_, β) = ν(λ_1_,γ) = ν(λ_2_,γ), ν_4_ = ν(λ_4_, α_0_) = ν(λ_4_, β) = ν(λ_4_,γ), and ν_5_ = ν(λ_5_, α_0_) = ν(λ_5_,β) =ν(λ_5_,γ).

The mutation probabilities for genealogy α_+_ are:

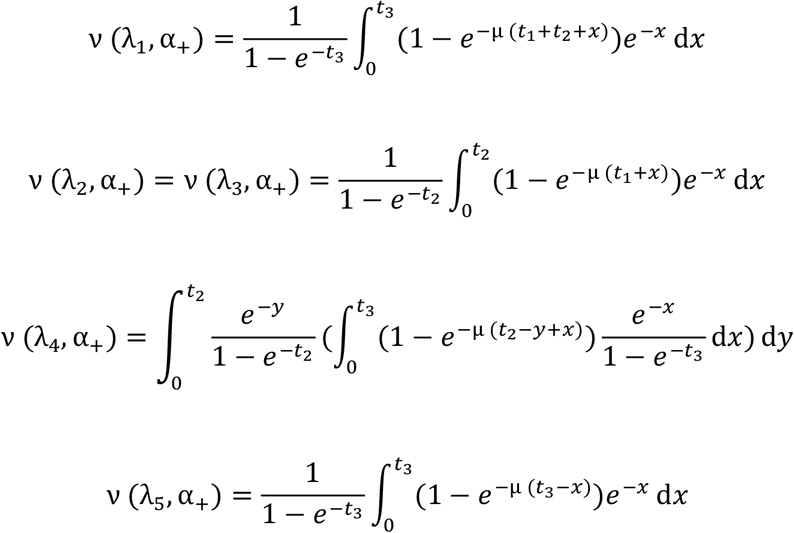

We use these functions, together with the probabilities of their corresponding genealogies (*p*(α_+_) = 1 − *e*^−*t*_2_^, and 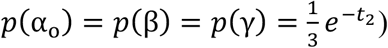, to obtain the values of *P_e_* and *P_o_* in equations 1 and 2.

### Calculating P_e_ / P_o_ for primates

To calculate *P_e_/P_o_* in the HC ancestor, we estimated branch lengths in coalescent units (6.8, 6.8, 7.6, 0.6, and 3.6 for H, C, G, HC and HCG, respectively) that agree with published divergence times and effective populations sizes (29, 30, 45). These lengths assume that H and C diverged 4.1 million years ago, that HC and G diverged 5.5 million years ago, and that generation time is 20 years throughout the clade. Additionally, we assume effective population sizes of 15, 15, 18, 19, 65 and 45 thousand individuals for H, C, G, HC and HCG, respectively. For the genome-wide average estimate of the population mutation rate, we use μ=3.6×10^−4^ per 30000 generations (i.e., with Ne = 15000 individuals and a per-site mutation rate of 1.2×10^−8^); the estimates for CpG and non-CpG sites used the appropriate per-site rates (31) and the same value of Ne. We assumed that the mutation rate was constant along the phylogeny.

### Calculating HRFs

An HRF (equation 3) can be calculated for any branch with a sibling, an ancestor, and two daughter lineages (cf. branch 4 in Figure 1). In other words, all branches of a phylogeny—except the root and leaves—have an HRF value. The *pepo* package walks up a phylogeny (in R, a *phylo* list as defined by the *ape* package; (46)) and calculates HRF for each internal branch, returning the values in a new data frame (compatible with *treeio*; (47)). The default HRF calculation in *pepo* allows reversals at the same rate as forward mutations, but these parameters can be specified by the user. This default method assumes that the ancestor of each focal branch has a length of at least 8N generations (i.e., *t*_3_=max(4, *x*), where *x* is the observed ancestral branch length). This setting, which can also be modified by users, is intended to reduce the effect of our assumption that A, B, and C have coalesced by the end of *t*_3_.

We calculated HRFs along the tomato phylogeny reported by Pease et al. (15), specifically the “best coalescent-based phylogeny from 100 replicates of MP-EST using 100 kb genome window trees” in the supplement of that paper. Because MP-EST assigns an arbitrary length of 9 to leaves, we modified terminal branch lengths in two ways. For species where multiple individuals were collected, we collapsed monophyletic samples into a single branch representing the species (the ancestral branch of the replicates). For species with a single sample, we assigned a terminal length of 1, an equally arbitrary value that is probably closer to the evolutionary history of this clade.

## Acknowledgements

This work was supported by National Science Foundation grant DBI-1564611.

---

**Figure S1.**
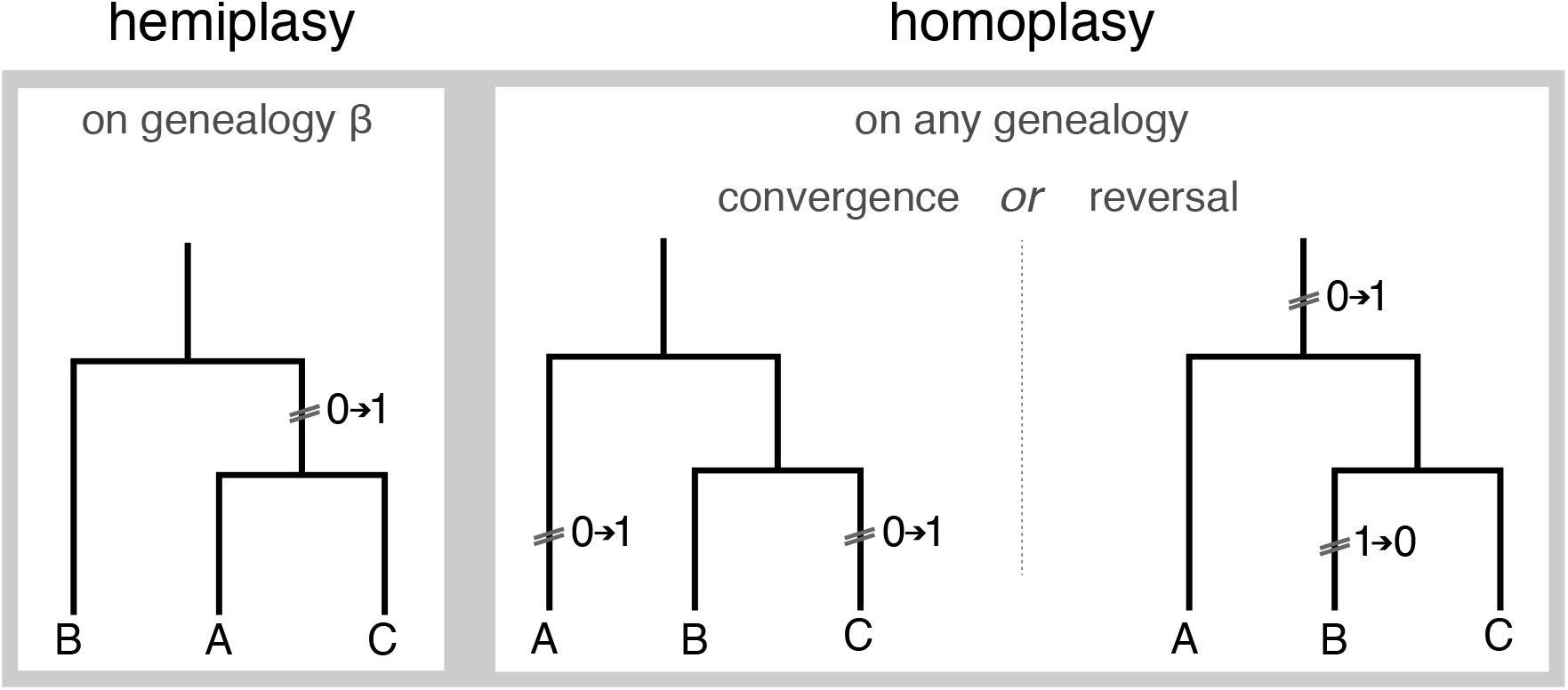
Diagram of possible scenarios of trait discordance.

**Figure S2.**
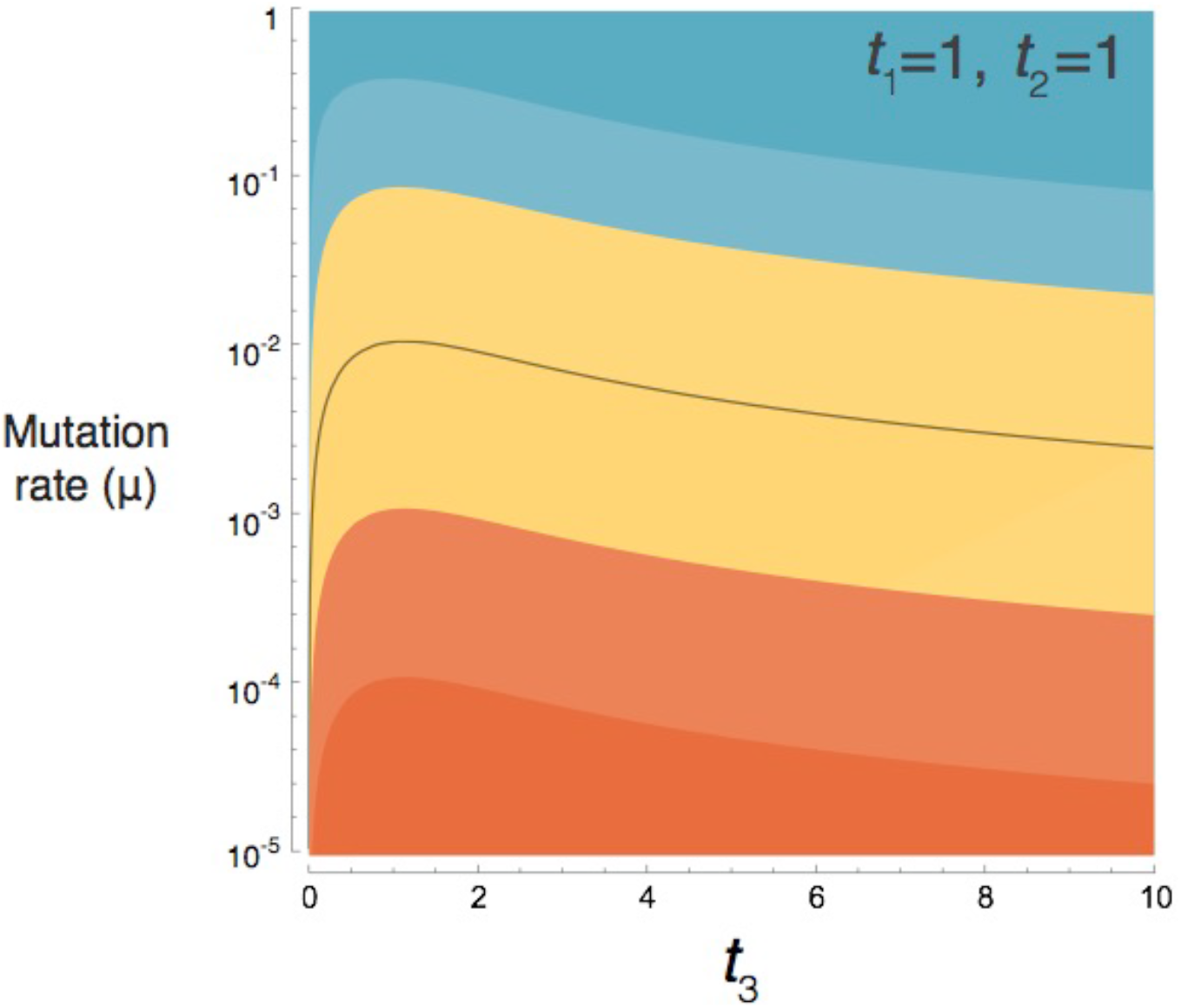
The relative probabilities of hemiplasy and homoplasy (*P_e_/P_o_*) are not dramatically affected by the length of the ancestral branch (*t*_3_; in units of 2*N* generations). Contours are as for Figure 2 (dark orange = hemiplasy more than 100x more likely than homoplasy). Rate of mutation shown along the vertical axis (μ, per 2*N* generations). The black solid line represents parameter values for which hemiplasy and homoplasy are exactly equal (*P_e_/P_o_* =1).

## References

1. Losos JB (2011) Convergence, adaptation, and constraint. Evolution 65:1827–1840.

2. Wake DB, Wake MH, & Specht CD (2011) Homoplasy: from detecting pattern to determining process and mechanism of evolution. Science 331:1032–1035.

3. Rosenblum EB, Parent CE, & Brandt EE (2014) The molecular basis of phenotypic convergence. Annual Review of Ecology, Evolution, and Systematics 45:203–226.

4. Storz JF (2016) Causes of molecular convergence and parallelism in protein evolution. Nature Reviews Genetics 17:239–250.

5. Duchêne S & Lanfear R (2015) Phylogenetic uncertainty can bias the number of evolutionary transitions estimated from ancestral state reconstruction methods. Journal of Experimental Zoology Part B: Molecular and Developmental Evolution 324:517–524.

6. Goldberg EE & Igić B (2008) On phylogenetic tests of irreversible evolution. Evolution 62:2727–2741.

7. Huelsenbeck JP, Nielsen R, & Bollback JP (2003) Stochastic mapping of morphological characters. Systematic Biology 52:131–158.

8. Maddison WP (2006) Confounding asymmetries in evolutionary diversification and character change. Evolution 60:1743–1746.

9. Pagel M, Meade A, & Barker D (2004) Bayesian estimation of ancestral character states on phylogenies. Systematic Biology 53:673–684.

10. Dunn CW, et al. (2008) Broad phylogenomic sampling improves resolution of the animal tree of life. Nature 452:745.

11. Misof B, et al. (2014) Phylogenomics resolves the timing and pattern of insect evolution. Science 346:763–767.

12. Beaulieu JM, O’meara BC, & Donoghue MJ (2013) Identifying hidden rate changes in the evolution of a binary morphological character: the evolution of plant habit in campanulid angiosperms. Systematic Biology 62:725–737.

13. Fontaine MC, et al. (2015) Extensive introgression in a malaria vector species complex revealed by phylogenomics. Science 347:1258524.

14. Jarvis ED, et al. (2014) Whole-genome analyses resolve early branches in the tree of life of modern birds. Science 346:1320–1331.

15. Pease JB, Haak DC, Hahn MW, & Moyle LC (2016) Phylogenomics reveals three sources of adaptive variation during a rapid radiation. PLoS Biology 14:e1002379.

16. Rokas A, Williams BL, King N, & Carroll SB (2003) Genome-scale approaches to resolving incongruence in molecular phylogenies. Nature 425:798–804.

17. Maddison WP (1997) Gene trees in species trees. Systematic Biology 46:523–536.

18. Hahn MW & Nakhleh L (2016) Irrational exuberance for resolved species trees. Evolution 70:7–17.

19. Avise JC & Robinson TJ (2008) Hemiplasy: a new term in the lexicon of phylogenetics. Systematic Biology 57:503–507.

20. Mendes FK & Hahn MW (2018) Why concatenation fails near the anomaly zone. Systematic Biology 67:158–169.

21. Robinson DF & Foulds LR (1981) Comparison of phylogenetic trees. Mathematical biosciences 53:131–147.

22. Copetti D, et al. (2017) Extensive gene tree discordance and hemiplasy shaped the genomes of North American columnar cacti. Proceedings of the National Academy of Sciences 114:12003–12008.

23. Mendes FK & Hahn MW (2016) Gene tree discordance causes apparent substitution rate variation. Systematic Biology 65:711–721.

24. Mendes FK, Hahn Y, & Hahn MW (2016) Gene tree discordance can generate patterns of diminishing convergence over time. Molecular Biology and Evolution 33:3299–3307.

25. Wu M, Kostyun JL, Hahn MW, & Moyle LC (2018) Dissecting the basis of novel trait evolution in a radiation with widespread phylogenetic discordance. Molecular Ecology

26. Hudson RR (1983) Testing the constant-rate neutral allele model with protein-sequence data. Evolution 37:203–217.

27. Pease JB & Hahn MW (2013) More accurate phylogenies inferred from low-recombination regions in the presence of incomplete lineage sorting. Evolution 67:2376–2384.

28. Scally A, et al. (2012) Insights into hominid evolution from the gorilla genome sequence. Nature 483:169–175.

29. Hobolth A, Christensen OF, Mailund T, & Schierup MH (2007) Genomic relationships and speciation times of human, chimpanzee, and gorilla inferred from a coalescent hidden Markov model. PLoS Genetics 3:e7.

30. Hobolth A, Dutheil JY, Hawks J, Schierup MH, & Mailund T (2011) Incomplete lineage sorting patterns among human, chimpanzee, and orangutan suggest recent orangutan speciation and widespread selection. Genome research 21:349–356.

31. Kong A, et al. (2012) Rate of *de novo* mutations and the importance of father’s age to disease risk. Nature 488:471–475.

32. Ané C, Larget B, Baum DA, Smith SD, & Rokas A (2007) Bayesian estimation of concordance among gene trees. Molecular Biology and Evolution 24:412–426.

33. Liu L, Yu L, & Edwards SV (2010) A maximum pseudo-likelihood approach for estimating species trees under the coalescent model. BMC Evolutionary Biology 10:302.

34. Lamichhaney S, et al. (2016) A beak size locus in Darwin’s finches facilitated character displacement during a drought. Science 352:470–474.

35. Li Y-Y, Liu Z, Qi F-Y, Zhou X, & Shi P (2016) Functional effects of a retained ancestral polymorphism in *Prestin*. Molecular Biology and Evolution 34:88–92.

36. Hahn MW, De Bie T, Stajich JE, Nguyen C, & Cristianini N (2005) Estimating the tempo and mode of gene family evolution from comparative genomic data. Genome research 15:1153–1160.

37. Suh A, Smeds L, & Ellegren H (2015) The dynamics of incomplete lineage sorting across the ancient adaptive radiation of neoavian birds. PLoS Biology 13:e1002224.

38. Zufall RA & Rausher MD (2004) Genetic changes associated with floral adaptation restrict future evolutionary potential. Nature 428:847–850.

39. Mallet J, Besansky N, & Hahn MW (2016) How reticulated are species? BioEssays 38:140–149.

40. Palesch D, et al. (2018) Sooty mangabey genome sequence reveals new insights into mechanisms of disease resistance in natural hosts of SIV. Nature 553:77–81.

41. Manceau M, Domingues VS, Linnen CR, Rosenblum EB, & Hoekstra HE (2010) Convergence in pigmentation at multiple levels: mutations, genes and function. Philosophical Transactions of the Royal Society B: Biological Sciences 365:2439–2450.

42. Projecto-Garcia J, et al. (2013) Repeated elevational transitions in hemoglobin function during the evolution of Andean hummingbirds. Proceedings of the National Academy of Sciences 110:20669–20674.

43. Zhen Y, Aardema ML, Medina EM, Schumer M, & Andolfatto P (2012) Parallel molecular evolution in an herbivore community. Science 337:1634–1637.

44. Lee KM & Coop G (2017) Distinguishing among modes of convergent adaptation using population genomic data. Genetics 207:1591–1619.

45. Prado-Martinez J, et al. (2013) Great ape genetic diversity and population history. Nature 499:471–475.

46. Paradis E, Claude J, & Strimmer K (2004) APE: analyses of phylogenetics and evolution in R language. Bioinformatics 20:289–290.

47. Yu G, Smith DK, Zhu H, Guan Y, & Lam TTY (2017) ggtree: an R package for visualization and annotation of phylogenetic trees with their covariates and other associated data. Methods in Ecology and Evolution 8:28–36.

